# Bilaterally Reduced Rolandic Beta Band Activity in Minor Stroke Patients

**DOI:** 10.1101/2021.10.15.464457

**Authors:** Joshua P. Kulasingham, Christian Brodbeck, Sheena Khan, Elisabeth B. Marsh, Jonathan Z. Simon

## Abstract

Stroke patients with hemiparesis display decreased beta band (13-25 Hz) rolandic activity, correlating to impaired motor function. However, patients without significant weakness, with small lesions far from sensorimotor cortex, nevertheless exhibit bilateral decreased motor dexterity and slowed reaction times. We investigate whether these minor stroke patients also display abnormal beta band activity.

Magnetoencephalographic (MEG) data were collected from nine minor stroke patients (NIHSS < 4) without significant hemiparesis, at ~1 and ~6 months postinfarct, and eight age-similar controls. Rolandic relative beta power during matching tasks and resting state, and Beta Event Related (De)Synchronization (ERD/ERS) during button press responses were analyzed.

Regardless of lesion location, patients had significantly reduced relative beta power and ERS compared to controls. Abnormalities persisted over visits, and were present in both ipsi- and contra-lesional hemispheres, consistent with bilateral impairments in motor dexterity and speed.

Minor stroke patients without severe weakness display reduced rolandic beta band activity in both hemispheres, which may be linked to bilaterally impaired dexterity and processing speed, implicating global connectivity dysfunction affecting sensorimotor cortex. Rolandic beta band activity may be a potential biomarker and treatment target, even for minor stroke patients with small lesions far from sensorimotor areas.

## 1 Introduction

Motor impairment is present in many stroke survivors (2), but does not always take the form of significant weakness. Patients with “minor stroke” and low National Institute of Health Stroke Scale (NIHSS) scores can exhibit normal strength but have disabling deficits manifesting as slowed response times and limited dexterity. This is common even in high functioning patients (3). Unlike hemiparesis, the underlying neural mechanisms for these processes are less well understood. These minor stroke patients also report difficulty with concentration and attention which, paired with decreased motor dexterity and slowed response times, hinder their ability to successfully return to work and reintegrate back into society. Previously, we found that such patients have low amplitude responses to visual stimuli that are temporally dispersed, possibly indicating a disruption of cortical networks (4). In this study we investigate neural responses in the sensorimotor cortex of the same cohort of minor stroke patients compared to age-similar controls, to determine if they display abnormal beta band activity, possibly linked to mechanisms underlying reduced motor dexterity and slowed response times.

Measurements of cortical activity using electroencephalography (EEG) or magnetoencephalography (MEG) indicate that rolandic beta band (13-25 Hz) responses are intricately linked to motor function (5–7). Spontaneous rolandic beta band activity may reflect multiple functional mechanisms in sensorimotor cortex including intracortical inhibition, communication, motor imagery and motor planning (8–10). Abnormal beta band activity has been observed in stroke (11,12), Parkinson’s disease (13) and other sensorimotor disorders (14). Stroke patients with motor deficits have been found to display reduced beta responses, especially in the ipsi-lesional hemisphere, possibly due to abnormal disinhibition and increased excitation (15). It is well established that beta band activity reduces during movement planning and execution (Event Related Desynchronization or ERD), and increases afterwards (Event Related Synchronization or ERS) in sensorimotor cortex (8,9,16). Although the neural mechanisms involved in these changes are not clear, prior work suggests that beta ERD may reflect cortical excitability and downregulation of inhibition while ERS may reflect active inhibition or a return to status quo after movement (17–19). Stroke patients with hemiparesis have decreased beta ERD/ERS, with a greater reduction in the ipsi-lesional hemisphere (11,20) and abnormal cortical patterns and latencies (21). However, it is unclear whether patients with small lesions without significant hemiparesis, would also display such abnormalities in beta band activity and beta ERD/ERS.

This study involved MEG data collected from stroke patients with small lesions with minor impairments in motor dexterity, and was motivated by several research questions. Firstly, we address whether even these minor stroke patients displayed abnormal rolandic beta activity compared to controls using relative beta power and beta ERD/ERS during button press responses. Next, we explore whether any abnormalities improve with time, using a subset of the patient cohort who returned for a second visit ~6 months later. Finally, we investigate whether the lesion location influences beta band activity by separately analyzing responses in ipsi- and contra-lesional hemispheres.

## 2 Methods

### 2.1 Subject Population

Nine patients with minor acute ischemic stroke 4-6 weeks postinfarct, and eight age-similar controls (age-matched within 5 years) without history of prior stroke were recruited for this MEG study (abnormalities in visual evoked responses from this population was reported in our previous study (4). The study was approved by the Johns Hopkins University institutional review board and all participants provided written informed consent. Patients with severe hemiplegia or aphasia, large vessel occlusions (M1 and M2 branches), prior history of dementia, incompletely treated psychiatric disease or uncorrected vision or hearing loss were excluded. Clinical scores NIHSS (22), Modified Rankin Scale (mRS; (23), Barthel Index (24) and cognitive tests Montreal Cognitive Assessment (MoCA; (25), and F-A-S verbal fluency test (26) were assessed. Along with clinical examination to assess strength, patients performed the grooved peg board test as a measure of motor speed and dexterity (27). Importantly, all patients exhibited low NIHSS scores and near normal neurological examinations with the exception of their cognition (measured using the MoCA-impaired defined as ≤26, see Table 1). Cognitive deficits were mild and improved between visits. Motor examinations were significant for mildly impaired rapid alternating movements contralateral to the side of the infarct for some (n=4), but uniformly slowed bilateral motor responses during task performance, and impaired dexterity as measured by the grooved pegboard. All patients displayed normal tone and strength to confrontation testing. Six patients and six controls returned for a 2^nd^ visit ~6 months postinfarct, before recruitment was halted due to the COVID-19 pandemic. A detailed description of patient characteristics is provided in Table 1. More details about the patient population, lesion locations and clinical measures are provided in our previous work (4).

**Table 1.** Clinical and Behavioral Measures. loc.: location of lesion S=subcortical, C=cortical, SC=subcortico-cortical; hemi.: lesion hemisphere; size: lesion volume in cc; GPBi: Grooved Peg Board ipsi-lesional hand reaction time (units in s.d. from the population average); GPBc: Grooved Peg Board contra-lesional hand reaction time. Patients show abnormal GPB scores bilaterally that improve but remain impaired for the 2^nd^ visit.

| Patient | Lesion |  |  | Acute | First visit |  |  |  |  | Second visit |  |  |  |  | Difference (Second - First) |  |  |  |  |
| --- | --- | --- | --- | --- | --- | --- | --- | --- | --- | --- | --- | --- | --- | --- | --- | --- | --- | --- | --- |
|  | loc. | hemi. | size | NIHSS | NIHSS | mRS | MoCA | GPBi | GPBc | NIHSS | mRS | MoCA | GPBi | GPBc | NIHSS | mRS | MoCA | GPBi | GPBc |
| 1 | SC | L | 10.5 | 4 | 0 | 1 | 24 | -1.5 | -3.6 | 0 | 1 | 26 | -2.8 | -0.6 | 0 | 0 | 2 | -1.3 | 3 |
| 2 | SC | L | 2 | 0 | 0 | 1 | 26 | -3.3 | -3.5 | 0 | 0 | 27 | -4.1 | -5.2 | 0 | -1 | 1 | -0.8 | -1.7 |
| 3 | SC | R | 1.6 | 3 | 3 | 1 | 21 | -5 | -14.9 | 0 | 1 | 20 | -8.4 | -14.1 | -3 | 0 | -1 | -3.4 | 0.8 |
| 4 | S | R | 0.4 | 2 | 2 | 1 | 24 | -15.7 | -14.1 | 0 | 0 | 29 | -0.4 | -1.1 | -2 | -1 | 5 | 15.3 | 13 |
| 5 | S | R | 0.3 | 1 | 0 | 0 | 28 | -7.8 | -9.3 | 1 | 1 | 30 | -3.2 | -12.1 | 1 | 1 | 2 | 4.6 | -2.8 |
| 6 | S | R | 6.8 | 3 | 0 | 1 | 28 | -2.1 | -6.1 | 0 | 0 | 27 | -0.8 | -4.5 | 0 | -1 | -1 | 1.3 | 1.6 |
| 7 | S | R | 1.3 | 12 | 0 | 1 | 28 | -0.9 | -1.7 |  |  |  |  |  |  |  |  |  |  |
| 8 | S | L | 2 | 4 | 1 | 2 | 22 | -4.5 | -3.2 |  |  |  |  |  |  |  |  |  |  |
| 9 | C | L | 0.7 | 1 | 0 | 1 | 27 | -14.8 | -1.9 |  |  |  |  |  |  |  |  |  |  |
| mean |  |  | 2.8 | 3.3 | 0.7 | 1 | 25.3 | -6.2 | -6.5 | 0.2 | 0.5 | 26.5 | -3.3 | -6.3 | -0.6 | -0.3 | 1.3 | 2.6 | 2.3 |
| SD |  |  | 3.5 | 3.5 | 1.1 | 0.5 | 2.7 | 5.6 | 5.1 | 0.4 | 0.5 | 3.5 | 2.9 | 5.6 | 1.5 | 0.8 | 2.3 | 6.8 | 5.6 |

### 2.2 Experiment Design

To investigate neural responses during motor activity, MEG data was collected while subjects performed picture-word matching tasks with button responses. A full description of each subtask is provided in our previous work (4). In brief, in each trial, subjects saw an image, followed 4 seconds later by a word and were asked to quickly and accurately press a button to indicate if the word corresponded to the image (yes: left button, no: right button). Reaction times were measured from the time the word first appeared on the screen to the time the participant pushed the response button. Across subtasks, subjects completed a total of 156 trials. Resting state data were also acquired in order to investigate if baseline beta band activity was different in patients versus controls. Resting state magnetic fields were recorded for ~1-2 minutes while the subjects rested with eyes open and fixated on a cross projected onto a screen ~2 feet in front of them.

### 2.3 MEG recording and preprocessing

MEG data was recorded using a 157 axial gradiometer whole head MEG system (Kanazawa Institute of Technology, Nonoichi, Ishikawa, Japan) while subjects rested in the supine position in a magnetically shielded room (VAC, Hanau, Germany). The data was recorded at a sampling rate of 1 kHz with a 200 Hz low pass filter, and a 60 Hz notch filter. All subsequent analyses were performed in mne-python (28,29), eelbrain (30) and R software (31). Saturating channels were excluded and the data was denoised using temporal signal space separation (32) to remove external noise. The MEG data was filtered from 1–40 Hz using an FIR filter (mne-python default settings), downsampled to 200 Hz, and independent component analysis was used to remove artifacts such as eye blinks, heartbeats, and muscle movements.

### 2.4 Neural source localization

The head shape of each subject was digitized using a Polhemus 3SPACE FASTRAK system, and head position was measured before and after the experiment using five marker coils. The marker coil locations and the digitized head shape were used to co-register the template FreeSurfer ‘fsaverage’ brain (33) using rotation, translation and uniform scaling. Single trial MEG data was source localized in the ‘ico-4’ surface source space, with current direction constrained to currents orthogonal to the white matter surface, using an inverse operator computed via Minimum Norm Estimation (MNE) (34), and a noise covariance estimated from empty room data. Neural sources in and around the pre- and post-central gyri were selected as the rolandic Region of Interest (ROI), using the ‘aparc’ parcellation labels ‘precentral’ and ‘postcentral’ (35); see Fig. 1A).

**Figure 1.**
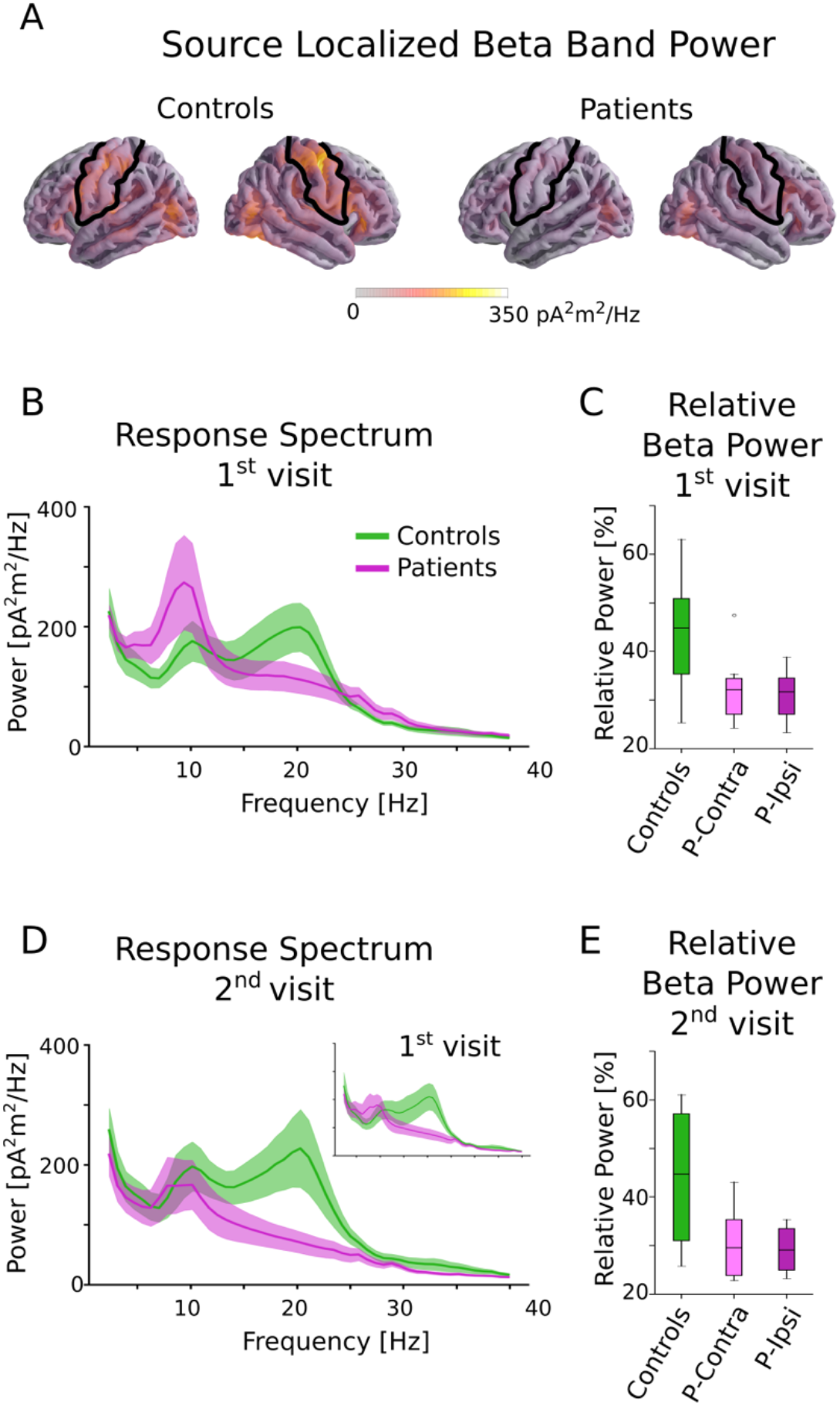
Beta power in controls and patients. **A**. Source localized beta band power for controls and patients averaged across all tasks. The black outline indicates the ROI used for all further analysis. Although patients and controls have similar beta activity in occipital areas, controls have much stronger activity in the rolandic ROI. **B**. Power spectral density of patients and controls averaged across all tasks in the central ROI for the 1^st^ visit. There is a clear group difference in the beta range (13-25 Hz). **C**. Relative beta power for controls and patients for the 1^st^ visit, separated by ipsi-lesional and contra-lesional hemispheres. Clear differences between controls and patients are seen, but there are no notable differences within patients between ipsi-lesional and contra-lesional hemispheres. **D**. Power spectral density for the 2^nd^ visit. Note that only a subset of subjects returned for the 2^nd^ visit. The 1^st^ visit spectrum averaged across only this subset of subjects is shown as an inset for comparison. **E**. Relative beta power for controls and patients for the 2^nd^ visit, separated by ipsi-lesional and contra-lesional hemispheres. Patients show reduced power even after ~6 months postinfarct. There are no notable differences within patients for ipsi- and contra-lesional hemispheres.

### 2.5 Frequency domain analysis

The source localized continuous MEG data during each picture-word matching task and the resting state was segregated into 15-second intervals. For each interval, the power spectral density was computed using Welch’s method (FFT length = 256 samples, with 50% overlap) and averaged across all segments and neural sources in the ROI. To account for individual variability in baseline frequency power, the relative beta power was computed by dividing the power in the beta range of 13-25 Hz by the total power in the range of 2-40 Hz. These relative beta powers were averaged across subtasks, log-transformed and were used for subsequent statistical analysis.

### 2.6 Event related (de-)synchronization analysis

The source localized MEG data in the rolandic ROI were epoched −3 to 3 seconds before and after the response button was pushed during the picture-word matching tasks. The time-frequency spectrograms for these trials were computed using Morlet wavelets (20 frequency bands with log-spacing in the range of 6-35 Hz with the number of cycles in each band being equal to half the center frequency). The beta ERD was computed as the percentage *decrease* in average beta (13-25 Hz) power in the time range of −1 s to 0.5 s relative to the button-press, compared to the baseline average beta power (time range −3 s to −2 s), consistent with established methods (9). The beta ERS was computed in a similar manner, as the percentage *increase* in average beta power over the baseline in the time range 0.5 s to 2.5 s. The trial averaged ERD/ERS values for each subject (averaged across subtasks) were used for subsequent statistical analysis.

### 2.7 Statistical analysis

The behavioral measures (reaction times and grooved peg board scores) were compared across patients and controls using t-tests with Bonferroni corrections for both the 1^st^ visit (9 patients) and the 2^nd^ visit (6 patients). Additionally, t-tests were used within the patient population to investigate if there were lateralization effects based on lesion hemisphere (ipsi- or contra-lesional sides). Correlations between behavioral and beta band measures were not investigated since there were no strong trends present during initial analysis.

To investigate group differences between patients and controls, the relative beta power in the rolandic ROI was analyzed with a linear mixed effects model using the ‘lme4’ package in R (36), since these models are capable of accounting for individual variation using random effects and for missing data (for this dataset, several subjects were not present for a 2^nd^ visit). This model was also used to test changes across visits, and across the picture-word matching tasks and resting state data. The dependent variable is relative beta log-power, with fixed effects of group (‘control’ or ‘patient’), task (‘matching’ or ‘resting’), and visit (‘1st’ or ‘2nd’) and a random intercept by subject. The full model with all interactions was tested for a significant difference over the reduced model without the highest-level interaction (group × task × visit) using the ‘drop1’ function in the lmerTest package in R, which performs a Type II ANOVA with the degrees of freedom estimated using Satterthwaite’s method (37). If the 3-way interaction was not significant, it was dropped from the model and the new model consisted of the 2-way interaction terms and the main effects. The same procedure with ‘drop1’ was used to check for significant interactions in this new model. If none of the 2-way interaction terms were significant, the final model only consisted of main effects.

To investigate possible hemispheric differences due to lesion hemisphere in the patient population, the beta log-power was separated by hemisphere, and paired two-tailed t-tests were used to test for a significant difference between the ipsi- and contra-lesional hemispheres. Separate t-tests with Bonferroni correction were performed for the 1^st^ visit (9 patients) and the 2^nd^ visit (6 patients).

A linear mixed effects model was also used to investigate group differences and changes across visits. This model had fixed effects of group, visit, and metric (‘ERD’, ‘ERS’) and random intercepts by subject. The same procedure as above was used to fit the models and determine significant interactions and effects, starting with the full model including the 3-way interaction. Finally, hemispheric differences in ERS and ERD due to lesion location were investigated within the patient population using paired two-tailed t-tests with Bonferroni correction for the 1^st^ and 2^nd^ visits similar to the above.

## 3 Results

### 3.1 Behavioral outcomes

Nine patients with minor stroke and eight age-similar controls without prior history of stroke were recruited for this study. Controls were recruited with similar age (within 5 years, mean [SD]; controls = 58 [13.1]; patients = 59.8 [15.7]) and sex (*n* male, controls = 4, patients = 4), but controls had a higher level of education that patients (years of education: controls = 18.6 [3.6]; patients = 14.1 [4.4]). Additionally, patients performed worse in cognitive tasks compared to controls (MoCA controls = 29.5 [3], patients = 26.0 [4], Mann-Whitney U Test p = 0.005). The clinical scores for all patients were low at both visits (mean [SD] NIHSS 1^st^ visit: 0.7 [1.1]; NIHSS 2^nd^ visit: 0.2 [0.4], mRS 1^st^ visit: 1 [0.5], 2^nd^ visit: 0.5 [0.5], see Table 1). Patients were bilaterally slow on the Grooved Peg Board task for both visits as shown in Table 1 (reaction time in units of SD from the mean, ipsi-lesional 1^st^ visit: −6.2 [ 5.6], 2^nd^ visit: −3.3 [2.9]; contra-lesional 1^st^ visit: −6.5 [5.1], 2^nd^ visit: −6.3 [5.6]). There were no significant differences between ipsi- and contra-lesional sides in the Grooved Peg Board task performance (1^st^ visit: t(8) = −0.15, p = 0.88; 2^nd^ visit: t(5) = −1.84, p = 0.12). The reaction times for the picture-word matching tasks were significantly longer for patients compared to controls for the first visit (mean [SD] Controls = 1.04 [0.44] s, vs. Patients = 2.17 [1.75] s; independent t-test on the reciprocal of the reaction times t(15) = 2.51, corrected p = 0.046, Cohen’s d = 1.3). Although reaction times improved (decreased) for both patients and controls for the 2^nd^ visit, there was no significant difference in the reaction time between the two visits (per subject difference between first and second visit: Controls = 62 ms [143 ms], Patients = −99 ms [318 ms]), and reaction times were still significantly different across groups for the 2^nd^ visit (t(10) = 2.72, corrected p = 0.04, Cohen’s d = 1.72). There was also no difference in reaction times for the ipsi- and contra-lesional button presses in patients, which were both impaired (1^st^ visit: t(8) = 0.88, p = 0.4; 2^nd^ visit: t(5) = −0.95, p = 0.39). Further details on reaction times for each subtask are provided in our previous work (4).

### 3.2 Beta power analysis

The MEG data during the picture-word matching tasks and resting state were source localized to the rolandic ROI and the (log-transformed) relative power in the beta frequency range of 13-25 Hz was computed (see Fig. 1). A linear mixed effects model with fixed effects group (‘patient’ or ‘control’), visit (‘1^st^’ or ‘2^nd^’) and task (‘matching’ or ‘resting’) and a random intercept by subject revealed a significant main effect of group (F(1,15) = 6.46, p = 0.023), but no other significant main effects (visit: F(1,39) = 0.65, p = 0.42; task: F(1,39) = 0.67, p = 0.42) or interactions (group × visit: F(1,36) = 0.06, p = 0.80; group × task: F(1,36) = 0.013, p = 0.91; visit × task: F(1,36) = 0.93, p = 0.34; group × visit × task: F(1,35) = 1.19, p = 0.28). A post-hoc t-test averaged over tasks and visits revealed that controls had significantly higher relative beta power than patients (t(15) = 2.54, p = 0.023, Cohen’s d = 1.31).

The effect of lesion hemisphere on the patients’ beta activity was investigated by separating the relative beta power into ipsi- and contra-lesional hemispheres. The relative beta power was not significantly different across the hemispheres for either the 1^st^ visit (picture-word matching: t(8) = 0.50, p = 0.63; resting: t(8) = 0.73, p = 0.49) or the 2^nd^ visit (picture-word matching: t(5) = 0.83, p = 0.44; resting: t(5) = 0.45, p = 0.67). Overall, patients had significantly reduced relative beta power for both the picture-word matching and resting tasks that did not improve for the 2^nd^ visit, and was independent of lesion hemisphere.

### 3.3 Event Related (De)Synchronization Analysis

Beta ERS and ERD were calculated for the button responses during the picture-word matching tasks using Morlet wavelet spectrograms (see Fig. 2; details in Methods). A linear mixed effects model for beta ERS/ERD with fixed effects group, visit, and metric (‘ERS’ or ‘ERD’) and a random intercept per subject was used to detect significant effects. The 3-way interaction was not significant (group × metric × visit F(1,36) = 2.02, p = 0.16). However, there was a significant interaction of group × metric (F(1,37) = 13.5, p < 0.001) as well as significant main effects of group (F(1,15) = 7.5, p = 0.015) and metric (F(1,40) = 22.7, p < 0.001). The other terms involving visit were not significant (group × visit: F(1,41) = 1.62, p = 0.21; visit × metric: F(1,37) = 0.34, p = 0.56; main effect of visit: F(1,46) = 0.04, p = 0.84). The significant interaction involving group and metric was analyzed further using post-hoc independent two-tailed t-tests with Bonferroni correction on the ERD and ERS averaged across visits and tasks. This revealed that patients had significantly lower ERS compared to controls (mean [SD] controls = 86.5 [44.3] %, patients = 39.9 [27.7] %, t(15) = 2.64, corrected p = 0.037, Cohen’s d = 1.36). Although the ERD also showed a similar trend, with controls being larger than patients, this difference was not significant (controls = 30.0 [9.4] %, patients = 24.8 [9.6] %, t(15) = 1.12, corrected p = 0.28).

**Figure 2.**
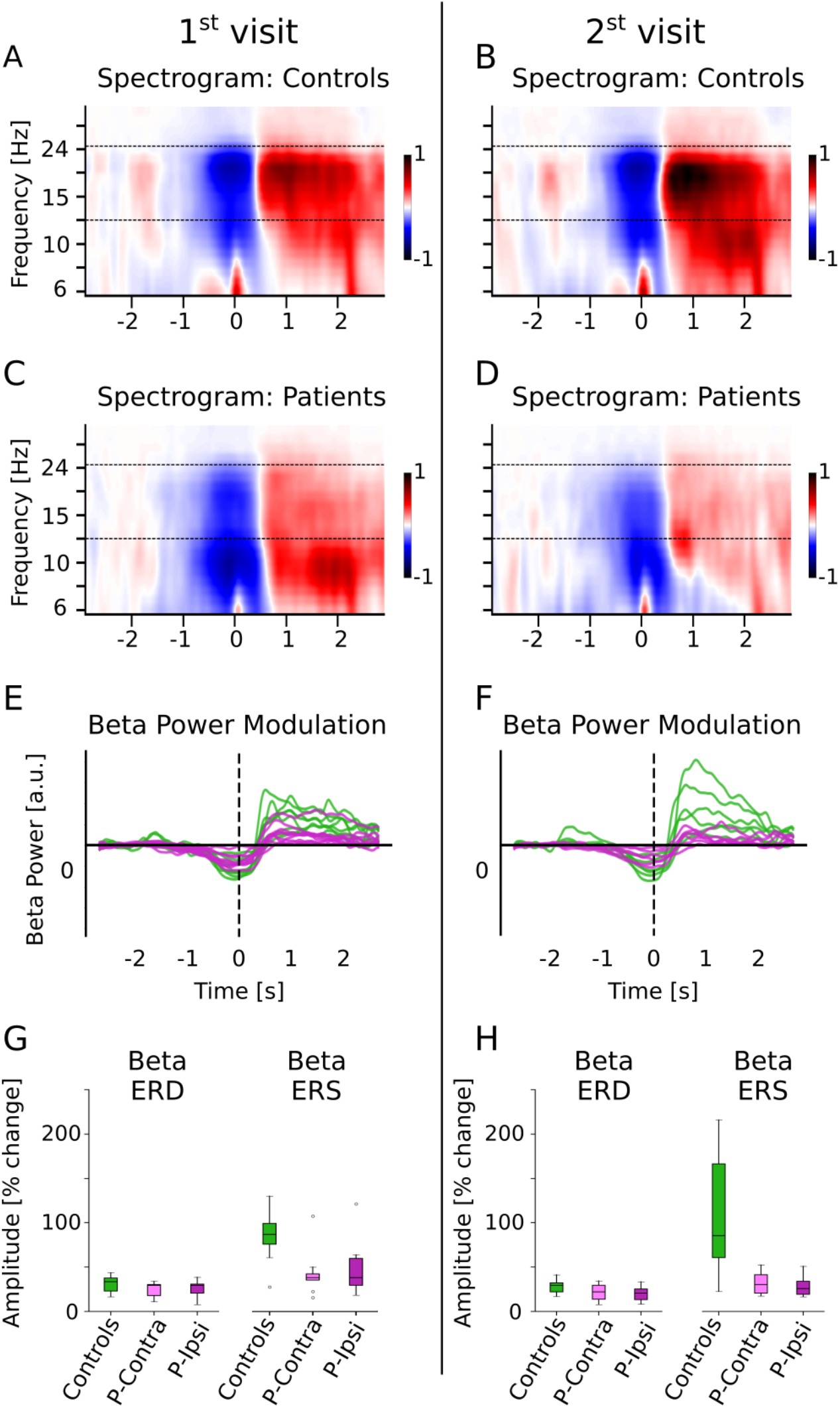
Beta ERD and ERS for controls and patients. **A, B, C, D**. The spectrograms for controls and patients (normalized w.r.t. baseline activity) are shown. The movement (button press) occurred at time t = 0. The dashed line indicates the beta band (13-25 Hz) that was used for further analysis. **E, F**. Beta power modulation per subject, computed as the average power in the beta band. Patients have reduced beta ERD/ERS. **G, H**. Beta ERD/ERS (% change from baseline) for controls and patients, separated by ipsi- and contra-lesional hemispheres. Patients have reduced ERD and ERS compared to controls for both visits, but show no differences across hemispheres. The group difference is much larger for the ERS than for the ERD.

To avoid the confound of ERS/ERD effects being driven by group differences during the baseline time-period (denominator in ERS/ERD calculations), the baseline beta power was calculated separately and was found to be not significantly different across groups (independent t-test t(15) = 1.0, p = 0.33). However, the beta power *relative* to total power in 2-40 Hz in the baseline time-period *was* significantly different between groups (independent t-test t(15) = 2.54, p = 0.022), in line with the reduction of spontaneous relative beta power in patients as shown in Fig. 1.

Finally, differences due to lesion location were tested by separating the ERD/ERS into ipsi- and contra-lesional hemispheres. There were no significant differences between the ipsi- and contra-lesional hemispheres within patients (ERD 1^st^ visit: t(8) = −0.21, p = 0.84; ERD 2^nd^ visit: t(5) = 1.62, p = 0.16; ERS 1^st^ visit: t(8) = −0.82, p = 0.43; ERS 2^nd^ visit: t(5) = 0.64, p = 0.55). Overall, patients had significantly lower ERS (but not ERD), in a manner that was consistent across tasks and visits and independent of lesion hemisphere.

## 4 Discussion

Minor stroke patients displayed reduced beta activity in rolandic areas compared to controls, both during picture-word matching tasks and resting state. Although these patients did not have significant motor impairment or hemiparesis, they reported difficulties with concentration, attention and an overall slowness combined with a reduction of motor dexterity, that hinders their ability to reintegrate well into society. We found that their rolandic beta activity was reduced bilaterally, independent of lesion hemisphere, and did not greatly improve with time. This is consistent with bilaterally slower reaction times and impaired grooved peg board scores that improved but remained below the normative average for their second visit. Interestingly, group differences in beta ERS were much larger than in beta ERD. A similar pattern of persistently reduced visual evoked responses was found in the same patient cohort in our previous study (4). Overall, these results suggest a more global disruption, not tied to lesion location, possibly involving long range cortical networks or a global excitation/inhibition imbalance, even for patients with only minor strokes.

### 4.1 Reduced beta activity in patients compared to controls

Minor stroke patients had significantly reduced relative beta power compared to controls in rolandic areas regardless of task. Although the beta ERD and ERS would be a significant component of the overall relative beta power during the picture-word matching tasks that involved button responses, we also found differences in relative beta power during the baseline period, which would not involve button press related motor beta activity. Spontaneous beta oscillations during resting were also reduced, consistent with prior work (15). This indicates that minor stroke patients have abnormal beta oscillations even without active movement.

### 4.2 Reduced beta ERS/ERD in patients compared to controls

Both beta ERS and ERD were reduced on average in minor stroke patients compared to controls, though only the ERS showed a significant difference. Prior studies have shown abnormal latencies, amplitudes and cortical patterns for both beta ERD and ERS in stroke patients with motor impairments (21,38–41). The beta ERD/ERS is thought to reflect regulation of intracortical inhibition (17), although they may also arise from several other functions including motor planning and short-term memory (9). The changes in beta ERD may depend on the specific motor pathology (39), and may not be large enough to detect in our population of minor stroke patients. A more thorough analysis with a larger population is required to ascertain whether the beta ERD also shows a significantly reduced amplitude, like the ERS, in minor stroke patients.

### 4.3 Beta abnormalities independent of lesion hemisphere

We found that for minor stroke patients, the reduction in beta activity was similar in both hemispheres, in contrast to some prior stroke studies that found a greater reduction of beta activity in the ipsi-lesional hemisphere (42,43). The observed bilateral reduction in beta band activity agrees with slower reaction times for both hands on the grooved peg board task in our patient population. Indeed, disinhibition and increased corticospinal excitability have been found in both hemispheres in cases of moderate to severe hemiparesis after unilateral stroke (44). Correspondingly, several studies do find bilateral abnormalities in beta activity due to stroke (12,44–46), consistent with bilaterally impaired hand dexterity after unilateral stroke (47). Several explanations for the mechanisms underlying these bilateral impairments have been proposed, including bilateral disinhibition, cortical reorganization and bilateral involvement in motor planning and execution (46,48). Additionally, these bilateral abnormalities may also indicate disruption of bilateral network connectivity even in areas without lesions (49–51).

### 4.4 Beta abnormalities are consistent across visits

Cognitive performance and reaction times for the minor stroke patients improved when they returned for their second visit, though remained abnormal compared to controls. Beta band measures showed little improvement between visits, unlike other studies which found that improved recovery from stroke correlates with a return to healthy levels of beta band activity (20,21,46). However, most of these studies involved patients with moderate to severe motor impairments, and may not be comparable to our population of patients with only minor motor deficiencies. In our case, patients still performed slower than average on the grooved peg board task even for the 2^nd^ visit, indicating a persistent reduction in dexterity despite some improvement. The changes in beta band measures during recovery from minor strokes may be quite small, and perhaps longer studies (>1 year postinfarct) are needed to detect such recovery effects. Alternatively, beta band abnormalities may persist due to mechanisms with long-term effects, such as network disruption (52).

In summary, we show that MEG beta band activity is a meaningful measure of motor deficits other than weakness in patients with minor stroke. The bilateral reduction in beta activity agrees with patients displaying an overall slowness and reduced dexterity in both hands and illustrates that even small infarcts may have bilateral impacts on motor function and sensorimotor activation, reflected in abnormal beta band activity. Prior studies indicate that these abnormalities are linked to a disruption of network connectivity or a global disinhibition of sensorimotor cortex, perhaps due to neural reorganization and compensatory mechanisms (44,53–55). This study has a limited number of subjects due to the need to stop recruitment during a pandemic. We could not disentangle the composite mechanisms causing this abnormal beta activity, nor could we detect neural correlates of motor performance and improvement with time, perhaps because of the small sample size. Future studies with larger populations of minor stroke patients and more complex motor task designs could shed light on the underlying causes of abnormal beta band activity. Nevertheless, it is important to note that even patients with small infarcts without weakness display such marked reductions in rolandic beta activity that agrees with impaired dexterity and reaction times, even in the contra-lesional hemisphere. Investigating whether these small lesions far from motor cortex can alter connectivity in remote areas could provide useful insights into the pathophysiology of these minor strokes and the mechanisms underlying bilateral motor impairments, and could lead to more effective methods of motor rehabilitation and recovery.

## Acknowledgements

This work was supported by an Innovative Research Grant through the American Heart Association (18IPA34170313) and National Science Foundation Grant SMA-1734892. This work is available online as a preprint (1).

## Notes

### Competing Interest Statement

The authors have declared no competing interest.

### Summary of Updates

More details on patient population. Minor changes to analysis and results. No conclusions were affected.

